# Possible changes in fidelity of DNA polymerase δ in ancestral mammals

**DOI:** 10.1101/2020.10.29.327619

**Authors:** Kazutaka Katoh, Naoyuki Iwabe, Takashi Miyata

## Abstract

DNA polymerase δ (polδ) is one of the major DNA polymerases that replicate chromosomal genomes in eukaryotes. Given the essential role of this protein, its phylogenetic tree was expected to reflect the relationship between taxa, like many other essential proteins. However, the tree of the catalytic subunit of polδ showed an unexpectedly strong heterogeneity among vertebrate lineages in evolutionary rate at the amino acid level, suggesting unusual amino acid substitutions specifically in the ancestral mammalian lineage. Structural and phylogenetic analyses were used to pinpoint where and when these amino acid substitutions occurred: around the 3′-5′ exonuclease domain in later mammal ancestry, after the split between monotremes and therians. The 3′-5′ exonuclease domain of this protein is known to have an impact on the fidelity of replication. Based on these observations, we explored the possibility that the amino acid substitutions we identified in polδ affected the mutation rate of entire chromosomal genomes in this time period.

## Introduction

DNA polymerase δ (polδ) is thought to synthesize the lagging strand during chromosomal replication in eukaryotes, while DNA polymerase ε (polε) is thought to synthesize the leading strand [1]. In each of polδ and polε, the catalytic subunit has two domains, the polymerase domain, which replicates DNA, and the 3′-5′ exonuclease domain, which proofreads errors made in the polymerase reaction. Interesting findings have recently been reported regarding these DNA polymerases, such as polδ’s cross-strand proofreading activity [2–4] and possible contribution of polδ to initiating replication of the leading strand [5, 6].

Since replication of chromosomes is essential for a cell, differences in the role of replicative DNA polymerases between eukaryotic taxa were rarely considered in functional studies, until recently as comparative analyses of these genes are becoming possible using sequence data from various organisms [4, 7, 8]. Here we report an unexpected observation based on a comparative analysis of the catalytic subunit of polδ, where lineage-specific rapid amino acid substitutions were found within quite a narrow organismal range, amniotes.

From an evolutionary viewpoint, polδ and polε, responsible for chromosomal replication, are of special interest; fidelity of these DNA polymerases can directly affect the mutation rate and the evolutionary rate of all genes on the chromosomal genomes, through germline replication. We also discuss this possibility.

## Results

### Polδ

Figure 1 shows an evolutionary tree based on the amino acid sequences of the catalytic subunit of polδ from vertebrates, using *Drosophila* and yeast as outgroups. For precise analysis, aves were excluded from this tree (see the “Avian polδ” subsection below for avian data). This tree shape is obviously strange in the mammalian linage (orange). Specifically, mammals in this tree have several times higher evolutionary rates at the amino acid level than reptiles (green) and other vertebrates. Also, the tree topology differs from the generally accepted one that the mammalian and reptilian lineages form sister-groups with each other [9]. The second point is most probably due to the well-known long-branch attraction (LBA) artifact [10] resulting from the first point. If assuming the sister-group relationship between mammals and reptiles, then the decrease of the log likelihood was only 9.78±11.8. Thus we focus on the first point, increase of the evolutionary rate, in the subsequent part of this report. Even if this tree topology is correct (ie, this tree contains paralogues or genes converged with paralogues), our discussion is unchanged in the point that mammalian polδ greatly differs from other vertebrates’ polδ.

**Figure 1:**
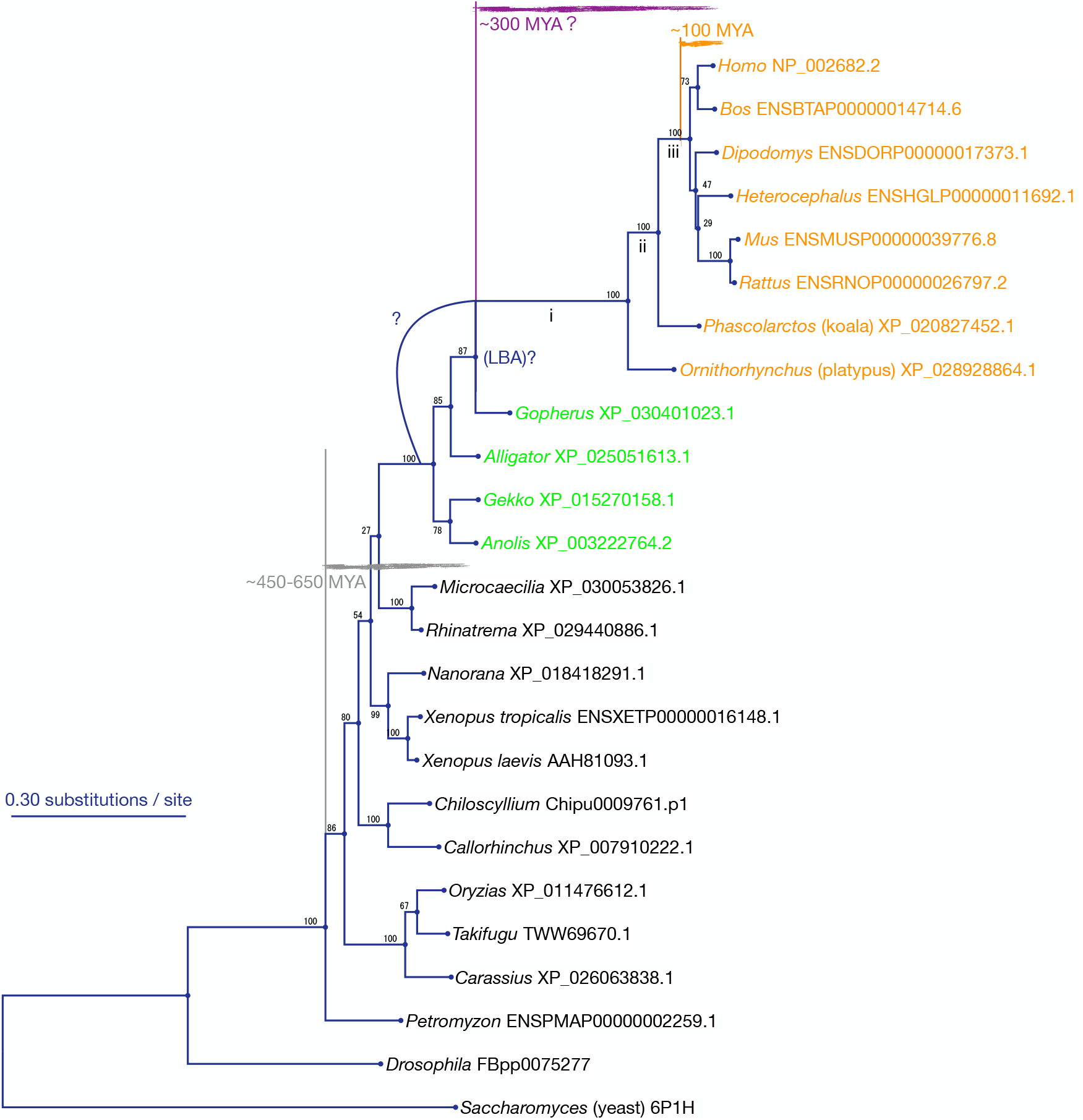
Evolutionary tree estimated using 910 amino acid residues of the catalytic subunit of polδ from vertebrates excluding aves. See Figure 4 for a tree including aves. The sequence data was taken from Ensembl and NCBI protein and manually sorted out to give an overview. i, ii and iii indicate time periods in ancestral mammalian lineage, defined in the main text. See Methods for calculation procedure.

When comparing purple and gray horizontal lines in Figure 1, the former is even longer in branch length despite the shorter elapsed time. Average evolutionary rate was estimated to be 1.40×10^−3^ substitutions/site/MYA and 2.84×10^−4^ - 4.10×10^−4^ subst./site/MYA for the purple (∼300 MYA) and gray (∼650 - ∼450 MYA) horizontal lines, respectively, where the approximate divergence times were taken from TimeTree [11]. Thus this tree is internally inconsistent in terms of timescale, or, in other words, has two or more different modes of evolutionary rate at the amino acid level. Note that this point holds even without the assumption of divergence times, as indicated by the tree shape.

This tree also implies that the strangely high evolutionary rate ended at some point in the mammalian lineage; After the radiation of eutherians (orange), the average evolutionary rate was estimated to be 6.06×10^−4^ - 9.32×10^−4^ subst./site/MYA (assuming the radiation of eutherians was 100 - 65 MYA), which are lower than that of ancestral+living mammals (purple; 1.40×10^−3^ subst./site/MYA). However, this estimation depends on the assumptions of divergence times, and is less clear than the above point.

### Polε

In contrast to polδ, polε resulted in an expected tree (Fig. 2), largely reproducing the accepted relationship between species. Difference in evolutionary rate between different lineages is not remarkable, unlike polδ.

**Figure 2:**
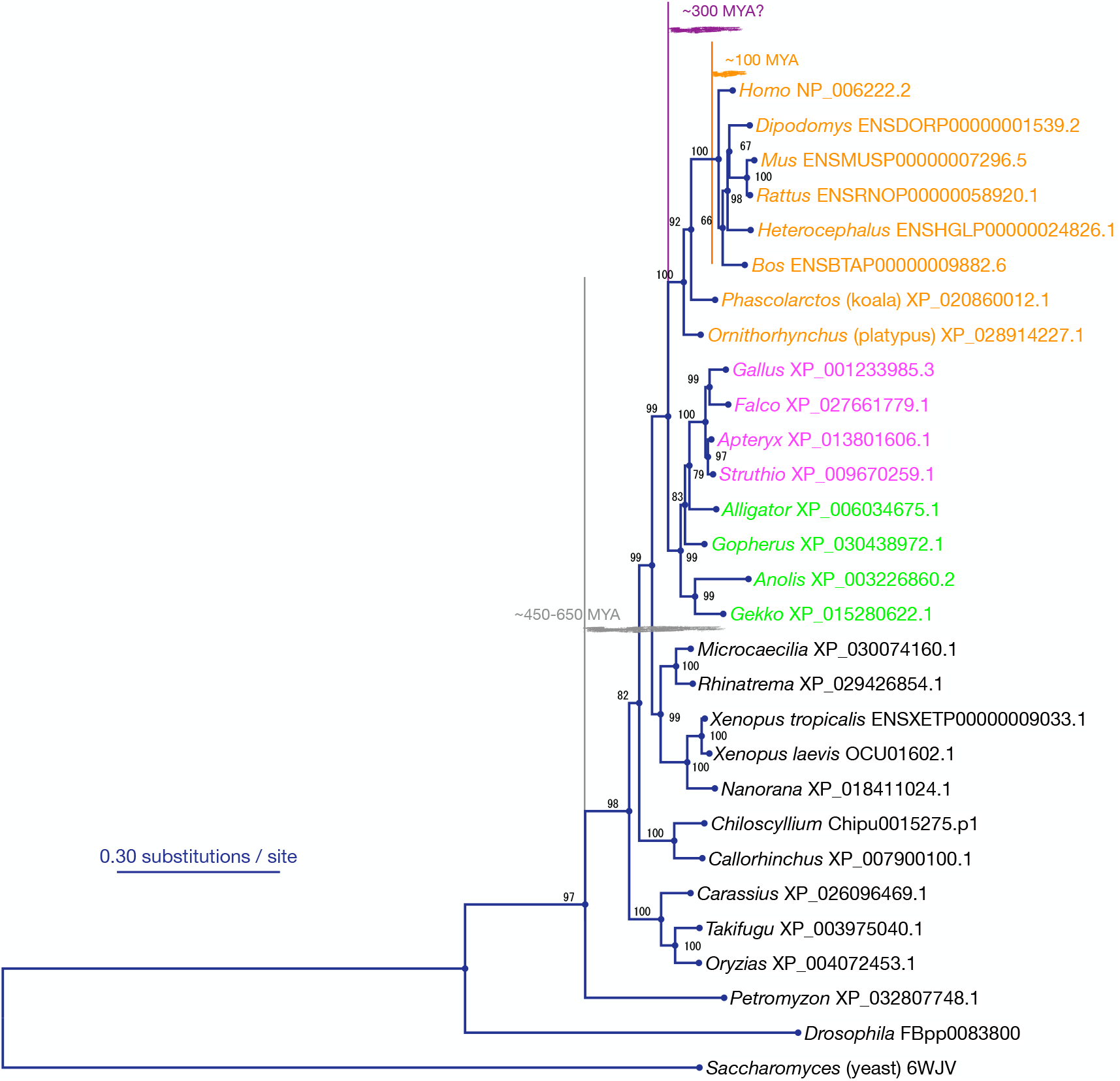
Evolutionary tree based on 1952 amino acid residues of the catalytic subunit of polε. See Methods for calculation procedure.

### Location of amino acid substitutions specifically in ancestral mammalian polδ

To narrow down the cause of the strangely high evolutionary rate at the amino acid level in polδ in the ancestral lineage of mammals, we explored which residues were substituted specifically in this lineage. First, we comprehensively collected amino acid sequences available from various large-scale sequencing projects, using aLeaves [12]. In total, 64 sequences from mammal, 20 sequences from reptiles and 123 sequences from other vertebrates were obtained, after manually excluding data with less information (short, redundant and/or having many ambiguous amino acids). On the multiple sequence alignment of this data, we checked the amino acid substitution patterns of 959 residues where structural information is available from yeast (PDB entry 6p1h). From this analysis we identified 16 amino acid residues that were substituted in the ancestral mammalian lineage (between the split of mammals from other vertebrates and the radiation of eutherians) but remained unchanged in each of non-mammalian vertebrates and eutherians. Their residue numbers in human (NP 002682.2 in NCBI reference sequence) and yeast (6p1h) sequences and the amino acids before and after each substitution are listed in Table 1. See the 16 rows that have a single amino acid in each of the second and third columns in this table.

**Table 1:** Amino acid residues that are conserved in each of eutherians and non-mammalian vertebrates and were substituted in the lineage of ancestral mammal. ^∗^, asterisk at the second column indicates that the substitution is across six physico-chemically different amino acid classes, ASGTP – MVIL – DENQ – C – KRH – YWF. **Residue numbers in bold** indicate that the residue is in the exonuclease domain (residues **309**–**538** in the numbering in yeast). ^†^, timing of substitution estimated using koala and platypus sequences. For definition and visual representation of periods i – iii, see main text and Figure 1. ^‡^, the correspondence of this residue between yeast and vertebrates is not obvious because of ambiguity of alignment. ←, the left arrow in the heading means that the direction of substitution can be estimated parsimoniously, as “non-mammalian vertebrates” form a paraphyletic group, not a monophyletic group. Residue number in 6p1h is based on the ATOM lines in the PDB file.

| Residue number in<br>NP_002682.2 | Amino acid in<br>eutherians | ←<br>Amino acid in<br>non-mammalian vertebrates | Timing of<br>substitution<sup>†</sup> | Residue number in<br>6p1h |
| --- | --- | --- | --- | --- |
| 127 | A * | M | ii | 139 |
| 164 | L * | F | i | 179 or 176 <sup>‡</sup> |
| 224 | R | K | i | 228 |
| **338** | C * | A/S | ii | **343** |
| **340** | L | M | i,iii | **345** |
| **341** | G * | V/M | ii | **346** |
| **355** | L * | F | i | **360** |
| **446** | T/S * | N | ii | **451** |
| **450** | S * | N | ii | **455** |
| **452** | V * | E | i,ii,iii | **457** |
| **525** | R | K | i | **530** |
| 528 | V * | C | i | 533 |
| 681 | L * | F | i | 688 |
| 829 | L | I | i | 835 |
| 831 | A | T | i | 837 |
| 847 | S * | C | i | 853 |
| 943 | L | I | i | 943 |
| 948 | H * | N | i | 948 |
| 1070 | I | L | iii | 1068 |

The 16 residues with lineage-specific changes were mapped onto the 3D structure of yeast homologue to test if the geometrical locations are significantly biased or not; the distribution of distances from the exonuclease catalytic center (green in Fig. 3; annotated in cd05777 in NCBI CDD) to these residues was compared the distribution of the distances from the same catalytic center to all residues in this protein. *p* value was estimated to be ∼0.023 by the Mann-Whitney U test under the null hypothesis that the two distributions are the same. On the other hand, the geometrical distribution of the same set of residues around the polymerase catalytic center (magenta; annotated in cd05533 in NCBI CDD) did not significantly differ from the distribution of all residues around the same catalytic center (*p*∼0.55).

**Figure 3:**
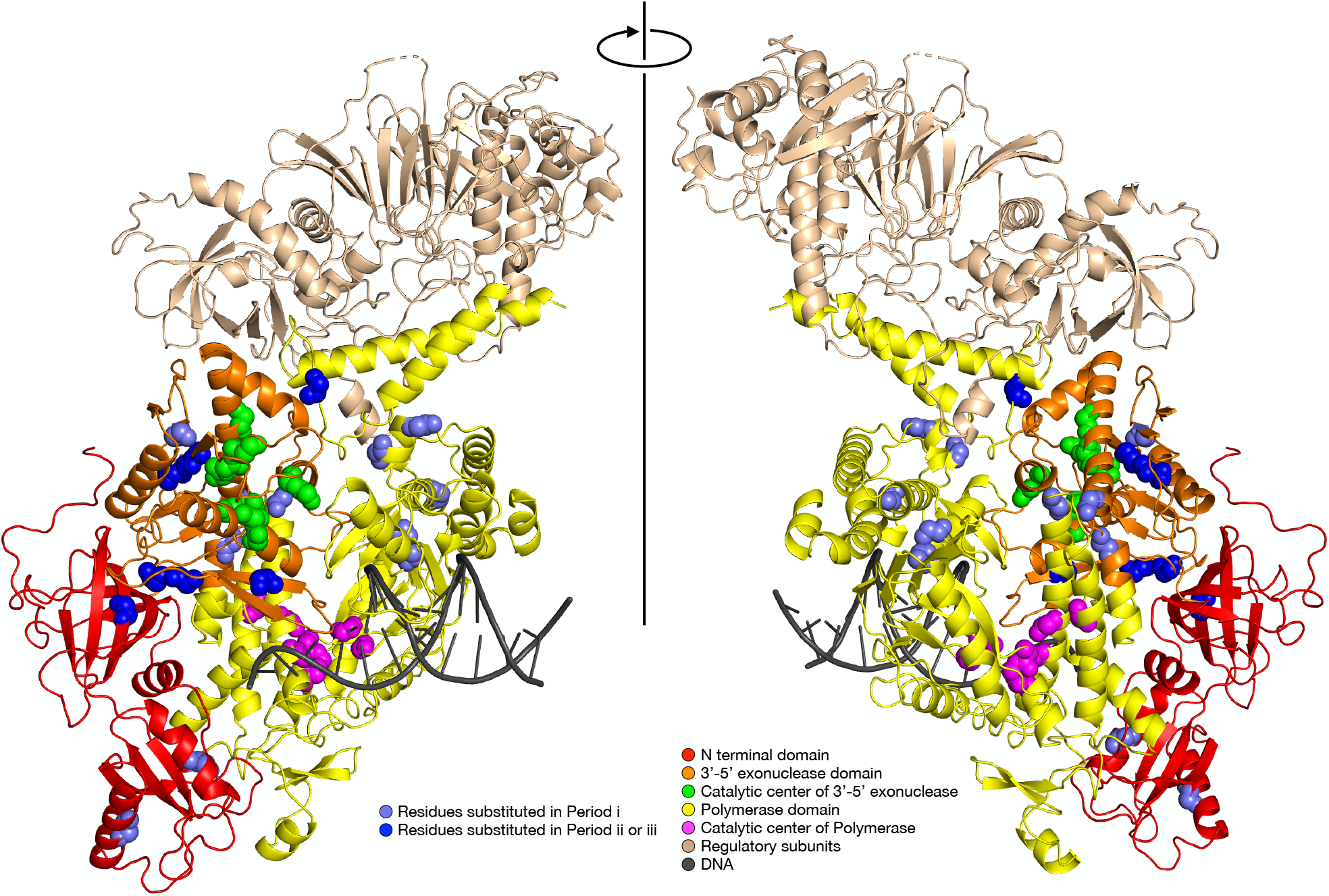
Mammal-specific amino acid substitutions mapped on 3D structure of polδ from yeast (6p1h). Slate blue, residues substituted in period i; Dark blue, residues substituted in period ii or iii (see main text and Figure 1 for definition of time periods).

To detect substitutions between physico-chemically dissimilar amino acids more clearly, we tried another criterion using “compressed amino acid alphabet”, where 20 amino acids are grouped into six classes (ASGTP – MVIL – DENQ – C – KRH – YWF) based on physico-chemical properties [13], and the changes across these six classes were counted. By means of this alphabet, 12 residues were found to be substituted specifically in the ancestral mammalian lineage (residues with ^∗^ in Table 1). This analysis resulted in a similar observation; the lineage-specific changes are distributed with a bias (*p*∼0.009) toward the exonuclease catalytic center.

### Timing of amino acid substitutions in ancestral mammalian polδ

The lineage-specific substitutions in ancestral mammal were further classified according to the timing when the substitutions occurred. By using marsupials and monotreme as key stones, the ancestral mammalian lineage can be divided into periods i – iii: Split between vertebrates/mammals — Period i — Split between monotremes/therians — Period ii — Split between marsupials/eutherians— Period iii — Radiation of eutherians. For visual representation of periods i – iii, see the branches indicated by i – iii in Figure 1. Table 1 shows the time (i, ii or iii) each substitution occurred, where the substitutions in later periods (ii and iii) clearly tend to be in the exonuclease domain (indicated by residue numbers in bold; 309–538 in the numbering in yeast). The locations of the mammal-specific substitutions on the 3D structure are graphically shown in Figure 3, colored slate blue (period i) and dark blue (period i or iii).

Our residue-wise analyses so far successfully narrowed down the location and the time period to be focused on: the 3′-5′ exonuclease domain, in a relatively short period in later mammal ancestry, ie, after the split between monotremes and therians, rather than immediately after the split between reptiles and mammals.

### Catalytic centers

All residues constituting the catalytic center of the exonuclease, however, are perfectly conserved in mammals, which is consistent with the exonuclease activity of mammalian polδ alive and having an impact on the fidelity of replication. The catalytic center of the polymerase is also fully conserved.

### Avian polδ

We also collected sequence data of avian polδ, which is very rare in databases and the reliability of sequences is questionable, being annotated as “low quality protein”. We obtained nearly full-length sequences of polδ only from three genus, *Falco*, *Melopsittacus* and *Catharus*, and fragmentary sequences from several other species. According to the phylogenetic tree including these sequences (Fig. 4), avian polδ had strange changes similar to (or more remarkable than) those found in mammals. First, the avian lineage (magenta) also has a high evolutionary rate at the amino acid level.

**Figure 4:**
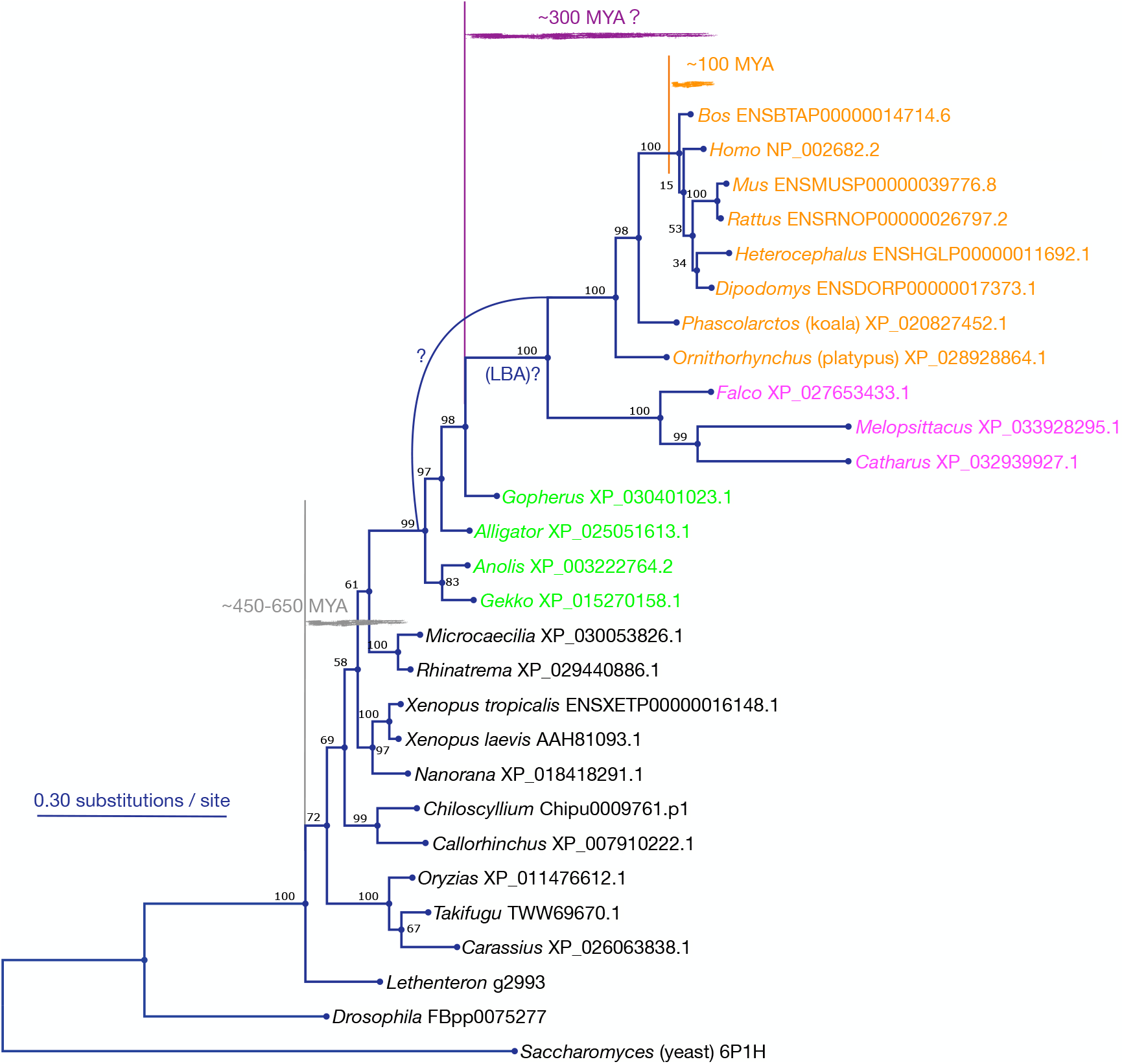
Evolutionary tree estimated using 824 amino acid residues of the catalytic subunit of polδ including avian data. *Falco* sequence was taken from NCBI Protein. To exclude the possibility of contamination or other artifacts of this sequence, we used three sequences (*Melopsittacus* and *Catharus* in addition to *Falco*), taken from RefSeq genomes, to confirm these sequences all result in similar trees.

The rate could be rather higher than that of mammals (orange), assuming no sequencing errors. Second, the phylogenetic relationship between amniotes (aves, mammals and reptiles) in this tree differs from the generally accepted one ((aves, reptiles), mammals) [9, 14, 15], and this tree topology is a typical one that was incorrectly estimated because of LBA. Even if this tree topology is correct (not affected by LBA), our discussion is unchanged; mammalian and avian polδ greatly differs from non-avian reptiles’ and other vertebrates’ polδ. These observations suggest that similar changes occurred in two independent lineages of homeotherms, mammals and aves. Since avian data is insufficient as noted in this subsection, we discuss, in the next section, the results obtained from mammalian data and just a few additional points derived from avian data.

## Discussion

We identified amino acid residues that were substituted specifically in ancestral mammalian polδ. These residues are located with a significant bias toward the 3′-5′ exonuclease domain. Further-more, the residues substituted in later mammal ancestry (periods ii and iii defined above) are quite clearly in/around the 3′-5′ exonuclease domain. This domain is involved in the proofreading function of polδ and known to strongly affect the fidelity of replication in both yeast and eutherians [16–18]. It is naturally deduced that these amino acid substitutions brought some drastic change in the fidelity of polδ. Then, was the change an increase or a decrease of fidelity? This question is also related to the evolutionary mechanism behind these amino acid substitutions; were they positively selected to increase or decrease the fidelity, or, were they a result of a relaxation of the functional constraints, ie, the fidelity of replication became simply less important?

To answer these questions, we focused on the special property of replicative DNA polymerases; their fidelity could affect the mutation rate of all genes on the chromosomal genomes, under an assumption that the replication mechanism of genomes is common between germline and somatic cells. It is known that the evolutionary rates at the DNA level of many genes (expectedly proportional to mutation rate) tend to be high in mammals than in other vertebrates. A useful index for mutation rate is synonymous substitution rate (*K_s_* [19] per divergence time) in orthologous genes, because *K_s_* is theoretically little affected by functional constraints on each gene and directly reflects mutation rate. In our analysis (see Methods) using the genes available in NCBI RefSeq genomes, *K_s_* between alligator and turtle was estimated to be around 0.4125 while *K_s_* between human (eutherian) and koala (marsupial) was estimated to be around 0.7875. The divergence times are thought to be ∼250 MYA and around ∼150 MYA for alligator/turtle and human/koala, respectively [11]. Thus the synonymous substitution rate (*K_s_* per divergence time) of the human/koala pair is about three times higher than that of the alligator/turtle pair. This is consistent with the possibility of the decrease of fidelity of polδ around the split between marsupials and eutherians.

These observations could be naively understood as follows. Fossil records indicate that ancestral mammals around the K-P boundary had small body size and short life span [20, 21], where the number of cell divisions and chromosome replications to reach reproductive maturity was small. Thus individuals with danger of frequent mutations (typically cancer prone) could have more chance to escape from purifying selection in such ancestral mammals than in longer-living vertebrates. As a result, the fidelity of DNA replication could be less important, ie, the balance between advantage and cost to keep the fidelity of replication could shift toward lower fidelity, in ancestral mammals. This scenario explains the strangely many amino acid changes around the proofreading (3′-5′ exonuclease) domain in this lineage around this period.

It is unclear if living eutherians still have a high mutation rate. The synonymous substitution rate between human and elephant (eutherians) was 0.2625/∼100 MYA, which is still about 1.5 times higher than that of alligator/turtle, 0.4125/∼250 MYA, but lower than human/koala, 0.7875/∼150 MYA.

The above interpretation also holds with the avian lineage; the evolutionary rate of aves is reportedly higher than that of the common ancestor of aves and non-avian reptiles [22–24]. This is partly consistent with the observation here; the avian lineage also had (or has) rapid amino acid changes in polδ, which possibly decreased the fidelity of replication. There are naturally other factors that affect evolutionary rates in different lineages; the increase of evolutionary rate at the DNA level in the avian and mammalian lineages is consistent with our observation, but the increase of the evolutionary rate in lepidosaurs and mouse [24] cannot be predicted from the amino acid substitutions in polδ.

Another notable feature of polδ is the GC richness (77%–90%) in the third codon position in aves, turtles, alligator and mammals. This range of taxa does not match the lineages with long branches (aves and mammals only) in the tree of polδ based on amino acid sequences (Fig. 4), but this lag might have a clue to understand the amino acid changes in avian polδ; we compared the GC content of the second codon position, strongly constrained by amino acid, of this gene between aves, turtles and alligator. In aves, the 5′ side (encoding the N terminal domain and exonuclease domain) of this gene is more GC rich (∼53%) than the 3′ side (∼45%) encoding the polymerase domain. Turtles and alligator (lineages with short branches in Fig. 4) have the opposite tendency: 5′ side, ∼46%; 3′ side, ∼51%. This difference might reflect the constraints at the amino acid level being weaken in the 5′ side of this gene in aves and a consequent shift toward GC at the DNA level because of the strong GC pressure that is common in aves, turtles and alligator.

The reduction of replication fidelity is, if any, not an exclusive cause of high mutation rate; short generation time and low replication fidelity can act cooperatively for high mutation rate (number of mutations per physical time) as discussed above. Therefore it is not easy to distinguish whether the high evolutionary rate at the DNA level is due to short generation time, low-fidelity replication, or both. A possible clue to distinguish these factors might be in a point that the mutations caused by polδ are expected to be biased toward the lagging strand in each round of replication. This point, strand-asymmetric mutation, has also been discussed in another context, a possible relationship to evolution at the morphological level [25, 26].

A further question would be whether reduction of fidelity in the replication system has any advantage or not, for mammals or homeotherms. Since they have high metabolic rate, a possible advantage might be a gain of replication speed by loose or relaxed proofreading.

It is certainly adaptive to keep the fidelity of replication to avoid too many mutations until an individual reaches reproductive maturity, while low-fidelity replication could have an advantage in special situations, like somatic hypermutation in B cells to effciently generate antibody diversity. Thus it can be speculated that the fidelity and speed of replication are regulated or tuned by using different combinations of subunits in the replication complex, at different developmental stages and/or in different organs. We tried to examine, based on the lineage-specific substitutions, a possibility that these amino acid substitutions formed a new interaction with an unrecognized subunit (eg, extrinsic 3′-5′ exonuclease) that works only in limited situations. However, we obtained no clear evidence for or against this hypothesis.

### Perspective

The hypotheses above cannot be proven just by in-sillico analyses and many other interpretations may be possible. To clarify what actually happened in ancestral mammalian polδ, experimental studies are critical and we believe the list of amino acid residues with lineage-specific substitutions (Table 1) is useful for this purpose. It is especially important and interesting to experimentally clarify the functional difference between the eutherian type of polδ, the ancestral type, being used by many other vertebrates, and intermediate types.

Our phylogenetic analysis including a small sample of avian polδ (Fig. 4) suggested the existence of avian-specific amino acid substitutions that might be more radical than mammalian-specific ones. This observation raises another interesting question; is there any link between the rapid amino acid substitutions in polδ and (seemingly subsequent or simultaneous) radiation and rapid diversification at the morphological level, in two independent lineages of homeotherms, mammals and aves?

## Materials and Methods

Amino acid sequence data was collected mainly using aLeaves [12], which has data from Ensembl and NCBI Protein. For avian polδ in Figure 4, RefSeq genomes were also used. Multiple sequence alignment was calculated using the L-INS-i option of MAFFT [27]. The alignment used for Table 1 is available at https://mafft.cbrc.jp/alignment/pub/pold/aln.oct5. Phylogenetic trees were estimated using RAxML [28] with the GAMMAIWAG model.

The number of synonymous substitutions (*K_s_*) between two species, *A* and *B*, was estimated as follows:

1. For each amino acid sequence in species *A*, selected the closest amino acid sequence in species *B* using BLAST [29]. Estimated *K_s_* for this gene pair using Yang and Nielsen’s method [30] implemented in PAML [31], regardless of whether this gene pair is orthologous or paralogous.
2. Drew a histogram of the distribution of *K_s_* with a bin width of 0.025. Used the leftmost (ie, with the lowest *K_s_*) peak to estimate *K_s_* between species *A* and *B*.

For a strict estimation, a set of orthologous pairs is necessary to estimate *K_s_* between two species. The procedure above approximates this step, based on an assumption that *K_s_* is roughly constant for every gene, since *K_s_* is little affected by functional constraints on each gene. Under this assumption, an orthologous gene pair between two species is expected to have a smaller *K_s_* value than paralogous pairs of the same gene between the same species pair, and thus the peak with the lowest *K_s_* was used for the estimation.

## Acknowledgements

This work was supported by JSPS KAKENHI 20K06767. We thank Mitsumasa Koyanagi, Martin C. Frith, Diego Diez, Songling Li, Kazuharu Misawa and Daron M. Standley for helpful discussions.

